# Genome sequence data of *Ascogaster reticulata* (Hymenoptera: Braconidae)

**DOI:** 10.64898/2026.10.06.756916

**Authors:** Kakeru Yokoi, Yooichi Kainoh, Miyuki Kanda, Seigo Kuwasaki, Kimura Kiyoshi, Hidemasa Bono

## Abstract

Parasitoid wasps develop in or on other arthropods and have evolved a wide range of strategies for exploiting their hosts. *Ascogaster reticulata* is a solitary egg–larval endoparasitoid wasp belonging to the subfamily Cheloninae of Braconidae. We constructed a genome assembly using long-read sequence data. The assembled genome was 280.5 Mbp, consisting of 209 contigs. The N50 value and GC content ratio were 13.9 Mbp and 30.43%, respectively. The percentage of transposable elements and repetitive regions was 52.43%. The predicted number of genes is 27,261. Merqury revealed that the QV value and completeness of the genome were 61.79 and 99.3917%, respectively. BUSCO analysis detected approximately 99% of the core genes. These results suggest that the genome data were of sufficient quality for use as a reference genome. Therefore, these data can be utilized as a genomic resource for comparative studies within Braconidae and for future investigations into the biology and evolution of Cheloninae parasitoid wasps.

## Introductions

Parasitoid wasps are a highly diverse group of insects that develop in or on other arthropods and have evolved a wide range of strategies to exploit their hosts. The family Braconidae (Hymenoptera) includes numerous parasitoid species that target different developmental stages in hosts and exhibit stage-specific modes of development. Such intimate associations require precise host recognition and coordination of parasitoid development with host growth and physiology^1,2^.

*Ascogaster reticulata* Watanabe is a solitary egg–larval endoparasitoid belonging to the subfamily Cheloninae of Braconidae (Figure 1). In Japan, it parasitizes tortricid moths, including the smaller tea tortrix *Adoxophyes honmai*. Females oviposit into host eggs, and the parasitoid develops internally as the host proceeds through larval development, eventually emerging from the host and pupating externally^3^. Female wasps locate and examine host egg masses during host-searching and oviposition behavior^4^. This prolonged association makes *A. reticulata* a useful system for studying parasitoid–host interactions. A previous study also suggested horizontal transfer of a mariner-like transposable element between *A. reticulata* and *A. honmai*^5^.

**Figure 1.**
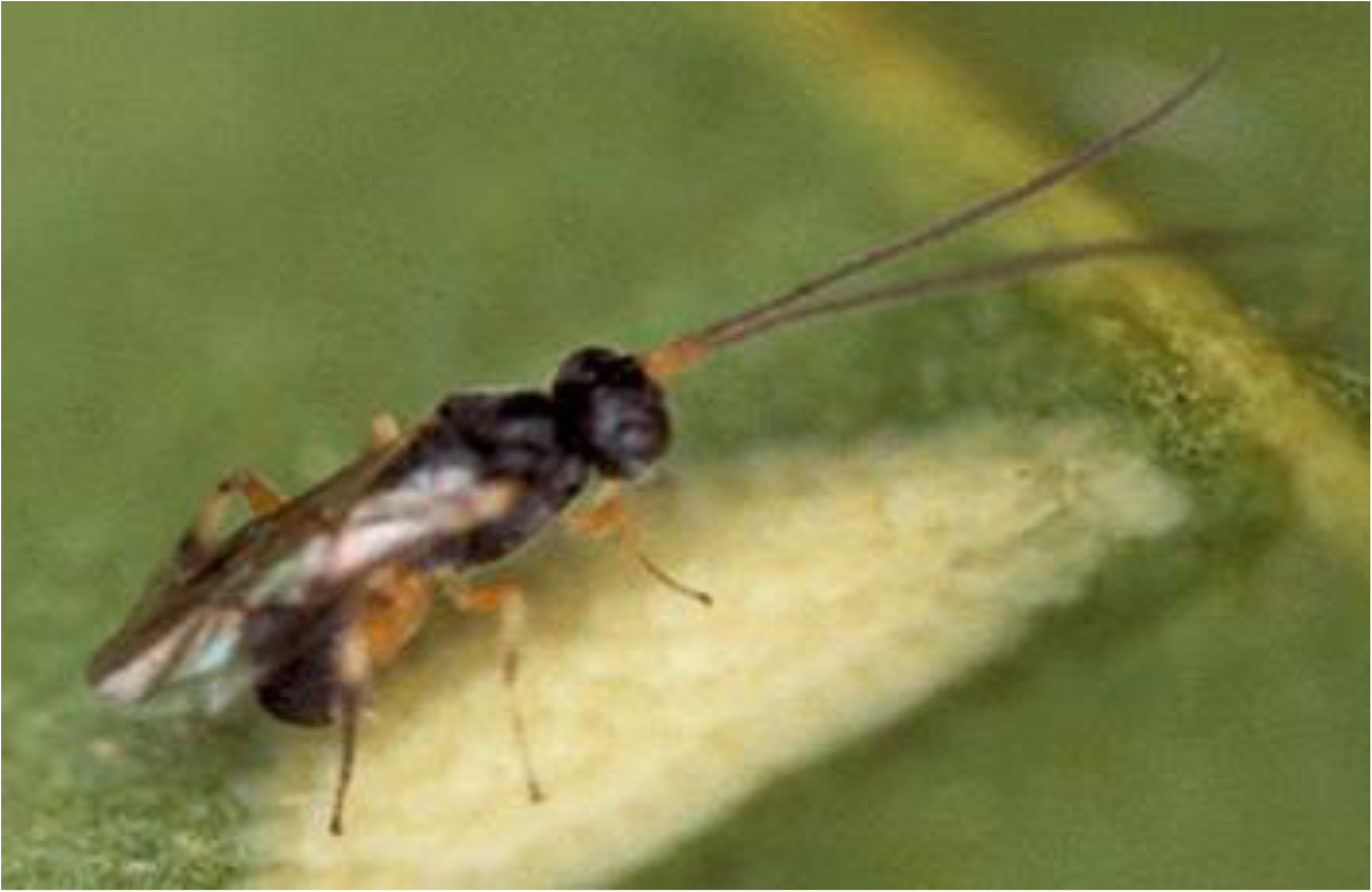
Picture of a female adult *A. reticulata*.

Genomic studies on parasitoid wasps in the family Braconidae have been conducted on several lineages, including members of the subfamily Microgastrinae, such as *Cotesia* and *Microplitis*, and Opiinae, such as *Fopius* and *Diachasma*^6–8^. These genomic resources have contributed to studies on genome evolution, parasitoid–host interactions, and the evolution of parasitism.

Genomic studies of several braconid wasps have revealed the importance of endogenous viral elements in parasitoid biology. In Microgastrinae, bracoviruses originate from viral sequences integrated into the wasp genome, and virus-derived particles deliver virulence genes that modify host immunity and physiology, thereby facilitating parasitoid development^7,9,10^. These studies demonstrate the value of genomic resources for understanding the diverse molecular mechanisms that have evolved in parasitoid wasps.

Genomic resources for the members of the Cheloninae subfamily remain comparatively limited. Genome assemblies have been reported for a small number of *Chelonus* species, including the chromosome-level assembly of *Chelonus formosanus*^11^, whereas genomic information for *Ascogaster* remains particularly scarce.

Cheloninae are characterized by an egg–larval parasitic strategy, in which parasitism begins during the host egg stage and parasitoid development continues as the host develops. This distinctive life history requires close coordination between parasitoid development and host physiology; however, its genomic basis remains poorly understood.

Genomic resources for Cheloninae are also relevant for understanding the diversity and evolutionary history of endogenous viral elements in Braconidae. Bracoviruses and other virus-derived genomic elements have been extensively investigated in some braconid lineages, particularly Microgastrinae. However, their occurrence, organization, and evolution vary among parasitoid lineages^10^. Expanding genomic resources for Cheloninae will therefore facilitate comparative analyses of genome organization, gene content, repetitive and virus-derived elements, and the evolution of parasitoid– host interactions within Braconidae.

In this study, we report the genome sequence of *A. reticulata* using HiFi long-read sequence data and provide its related data including gene set data and its functional annotation data (Figure 2). Also, we performed several quality assessments of the genome data of *A. reticulata*.

**Figure 2.**
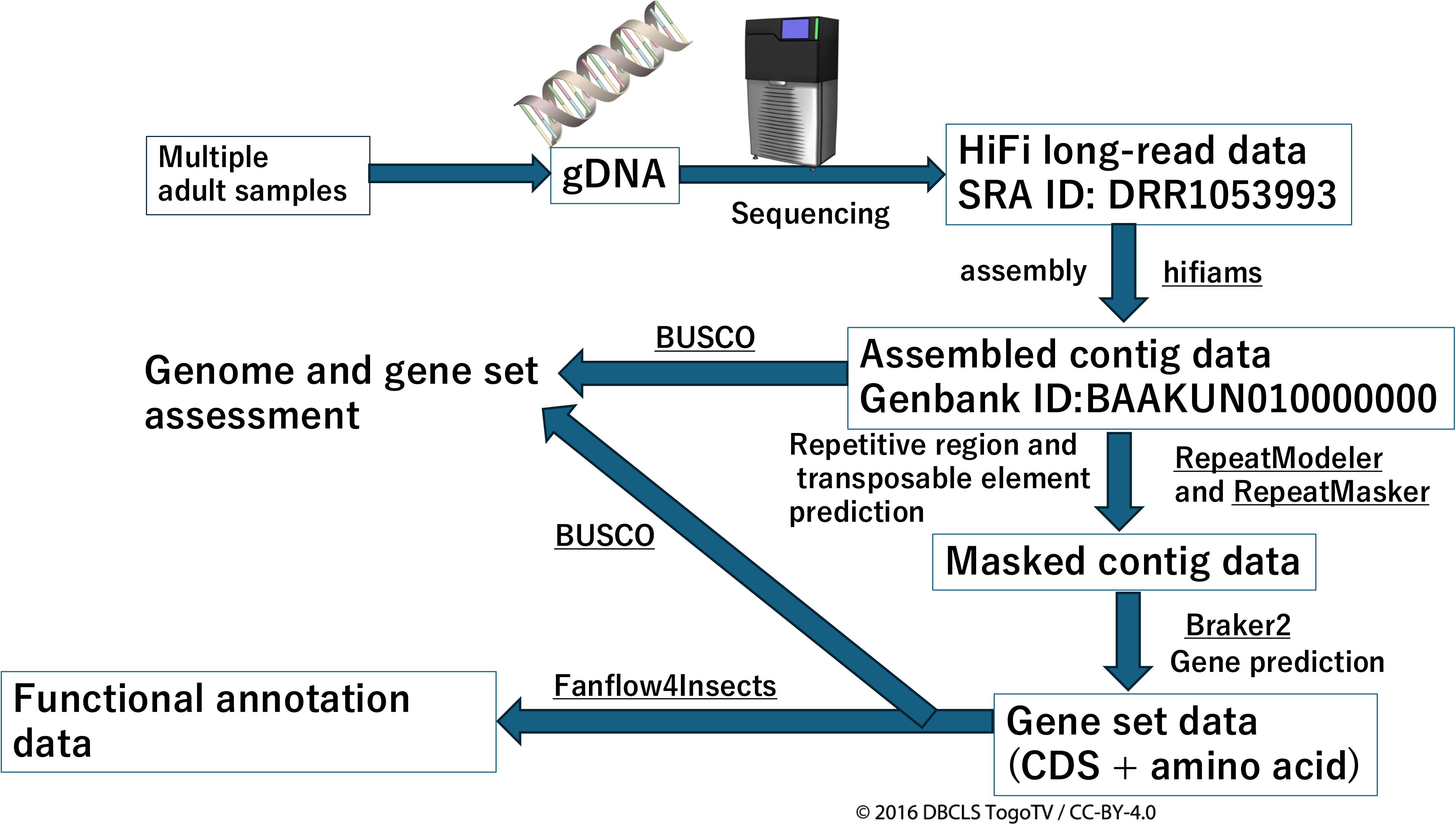
Workflow of data analysis in this study. Samples or genome and data names are shown in boxes. Data, except masked genome data in the figure, are available in the public database or Figshare (see “Data Availability” section), and codes or scripts used during data analysis in this study can also be accessed from Figshare (see “Code Availability” section). The software names are underlined. The images of genome DNA and the sequencer are from TogoTV (© 2016 DBCLS TogoTV, CC-BY-4.0 https://creativecommons.org/licenses/by/4.0/.“).

## Materials and Methods

### Sample preparation

The parasitoid *A. reticulata* was reared as described by Kainoh (1988)^12^, and its host *A. honmai* was reared as described by Tamaki (1966)^13^. Colonies of parasitoids and their hosts were obtained from stock cultures reared under laboratory condition (25 ± 1 °C, 60 ± 10% RH, and L16:D8 photoperiod). Parasitized and unparasitized *A. honmai* egg masses deposited on the wax paper were sterilized with 70% ethanol, washed with distilled water, and inoculated into plastic boxes (25 × 18 × 8 cm) containing an artificial diet and crumpled pieces of wax paper (9 × 4.5 cm). The hatched larvae were reared in these boxes for 18 days after inoculation. *A. reticulata* cocoons and *A. honmai* pupae were collected from rearing boxes. The cocoons were individually placed in test tubes (1 cm in diameter, 7.5 cm in length, 5 mL) with cotton plugs until they emerged. *A. reticulata* adults were separated by sex, and 10–20 of each were kept in a plastic container (15 cm in diameter, 9 cm in height) lined with moist cotton and honey on the inner wall. Two- to four-day-old naive female wasps were used for the colony rearing. *A. honmai* pupae were separated by sex and placed in rearing boxes (25 × 18 × 8 cm) with moist cotton. After emergence, male and female moths were mixed in boxes, and the inner side of the box lid was covered with a sheet of wax paper (37 × 25 cm) for oviposition. The egg masses were exposed to adult parasitoids, and the resulting offspring were used to establish the next generation. Adult male parasitoids 2–4 d old were used in the experiments. The ten adult males were pooled to obtain a sufficient amount of genomic DNA. The laboratory colony was considered to have relatively low genetic diversity among individuals because of prolonged inbreeding. For this reason, the pooled samples were used for HiFi genome sequencing, despite the general recommendation to use DNA from a single individual.

### Genomic DNA extraction and sequencing

Genomic DNA of *A. reticulata* was extracted from 10 adult males using a Nanobind PanDNA kit (Pacific Biosciences, CA, USA). The genomic DNA was fragmented to a target size of approximately 12 kb using the g-TUBE (Covaris, Woburn, MA, USA). The construction of SMRTbell libraries was accomplished using the SMRTbell Prep Kit 3.0, following the manufacturer’s protocol. Single-molecule sequencing was performed using the Sequel IIe system (Pacific Biosciences, CA, USA).

### Data analysis

The data analysis workflow is shown in Figure 2. Genome sequences were constructed from raw HiFi sequence data using Hifiasm (version v0.19.8-r603)^14–16^. Basic statistics of the constructed genome, such as total length and N50, were calculated using Assembly-Stats (version 1.0.1)^17^. The assembled genome data were assessed using Merqury (version 1.3)^18^ and BUSCO (version 5.2.2) against the Insecta_odb10 database^19^. To detect repeat regions and TEs in the constructed genome, RepeatModeler2 (version DEV) and RepeatMasker (version 4.1.2-p1) were used^20,21^. The regions (sequences) of the predicted repeats and TEs were replaced with lowercase characters that were not used for gene prediction, as described below, and referred to as “masked genome sequence data.” Using the masked genome sequence data as input, gene prediction was performed using Braker2 (version 2.1.6) with the Arthropod OrthoDB protein file (v10) from OrthoDB^22,23^. To perform gene functional annotation, the predicted amino acid sequences were used as input data against reference protein datasets from model organisms, which included *Drosophila melanogaster* (version Release 6 plus ISO1 MT; Drosophila_melanogaster.BDGP6.32.pep.all.fa.gz)^24^, *Caenorhabditis elegans* (version WBcel235; Caenorhabditis_elegans.WBcel235.pep.all.fa)^25^, Homo sapiens (version GRCh38; Homo_sapiens.GRCh38.pep.all.fa.gz)^26^, and *Mus musculus* (version GRCm39; Mus_musculus.GRCm39.pep.all.fa.gz)^27^. Additionally, they were also searched against Unigene (Uniprot KB protein data, uniprot_sprot.fasta.gz)^28^, Pfam (Pfam-A.hmm)^29^. The annotation was performed using Fanflow4Insects^30^.

### Data records and Code Availability

Long-read sequencing data were deposited in the DNA Data Bank of Japan (DDBJ) Sequence Read Archive (SRA) (ID: DRR1053993)^31^. The assembled genome data are available in NCBI GenBank (Accession ID: BAAKUN01000000)^32^. The list of contig IDs by Hifiasm and accession IDs in GenBank^33^, RepeatModeler output data (TE consensus sequence), RepeatMasker output data (gff, tbl, and out files)^34^, predicted gene set data (gff file, cDNA sequence fasta file, and amino acid sequence fasta file)^35^, and functional gene annotation data^36^ can be obtained via Figshare. All supplemental data are accessible in Figshare via a single URL^37^. All scripts used in this study are described in txt files available in Figshare^38^.

## Results and Discussions

The *A. reticulata* genome sequence data were constructed as shown in Figure 2.The total length of the assembled genome was 280,531,855 bp (genome coverage was approximately 165x), consisting of 209 contigs. The average contig length and size of the largest contig in the genome sequence data were 1,342,257.68 bp and 24,055,980 bp, respectively (Table 1). The N50 was 13,935,451 bp, which was sufficient for reference genome data. The percentage of GC content was 30.43%, and the percentage of transposable elements (TEs) and repetitive regions in this genome was 52.43%, of which sequences in the genome were masked and not used for gene prediction. The number of TEs and repetitive sequences are summarized in Table 2. DNA elements (Class II) were the most enriched TEs in this genome, except for unclassified TEs. Fewer class I TEs (SINEs, LINEs, and LTR elements) were identified than class II TEs. Repetitive regions (satellites, simple repeats, and low complexity) comprised approximately 16.2% of the *A. reticulata* genome. Using the masked sequence data (masked sequence is not used for gene prediction.), 27,261 genes were predicted, and the functional annotation was performed.

**Table 1.** Basic status of the assembled *A. reticulata* genome

|  |  |
| --- | --- |
| Total length | 280,531,855 bp |
| Number of contigs | 209 |
| coverage | 165 |
| Average value of the contig length | 1,342,257.68 bp |
| Size of the largest contig | 24,055,980 bp |
| N50 | 13,935,451 bp |
| GC percent | 30.43% |
| Percent of transposable elements or repetitive regions | 52.43% |
| Number of predicted genes | 27,261 |

**Table 2.** Transposable elements and repetitive region in *A. reticulata* genome

| Category | number of elements | length of occupied | percentage of sequence |
| --- | --- | --- | --- |
| SINEs | 678 | 197,829 bp | 0.07% |
| LINEs | 5,110 | 7,243,791 bp | 2.58% |
| LTR elements | 8,505 | 10,110,787 bp | 3.60% |
| DNA elements | 30,248 | 24,685,761 bp | 8.80% |
| Unclassified | 149991 | 58,772,411 bp | 20.95% |
| Small RNA | 1164 | 504,354 bp | 0.18% |
| Satellites | 189 | 31,021 bp | 0.01% |
| Simple repeats | 148,119 | 43,234,760 bp | 15.41% |
| Low complexity | 27708 | 1,868,276 bp | 0.67% |

Multiple assessments were performed to determine whether the constructed genome data were of sufficient quality to serve as a reference genome. Merqury analysis revealed that the QV value and completeness of the assembled genome were 61.79 and 99.3917%, respectively, based on the k-mer frequency of HiFi reads. The copy number spectrum obtained using Merqury (k=21) is shown in Figure 3. The distinct red peak at approximately kmer_multiplicity = 105 indicates that the majority of k-mers existed as a single copy in the haploid assembly, demonstrating a lack of artificial duplications. The gray peak near the origin represents low-frequency k-mers found only in the sequencing reads, corresponding to sequencing errors that were correctly filtered during assembly. BUSCO analysis shows the completeness of the constructed genome or predicted gene set by searching core genes (named “BUSCOs”) of insects in the database “insecta_odb10.” We performed BUSCO analysis using the genome sequence as a query, where 99% of the core genes were detected (98.7% as single copies), whereas only 1% of the core genes were fragmented or undetected (Table 3). As described above, the N50 value of the genome data was approximately 14 Mbp, meaning that this genome data possesses high contiguity (Table 1). In the NCBI Genome Data Viewer, 14 Braconidae genome datasets were registered (https://www.ncbi.nlm.nih.gov/gdv?org=cotesia-typhae&group=braconidae) as of 2026/8/1. The genome sizes of the Braconidae insects range from 106.7 to 384.4 Mb (Table 4). The size of the *A. reticulata* genome (280.5 Mb) was comparable to that of Braconidae insects. These results suggest that the assembled *A. reticulata* genome data possess sufficient quality for use as a reference genome.

**Figure 3.**
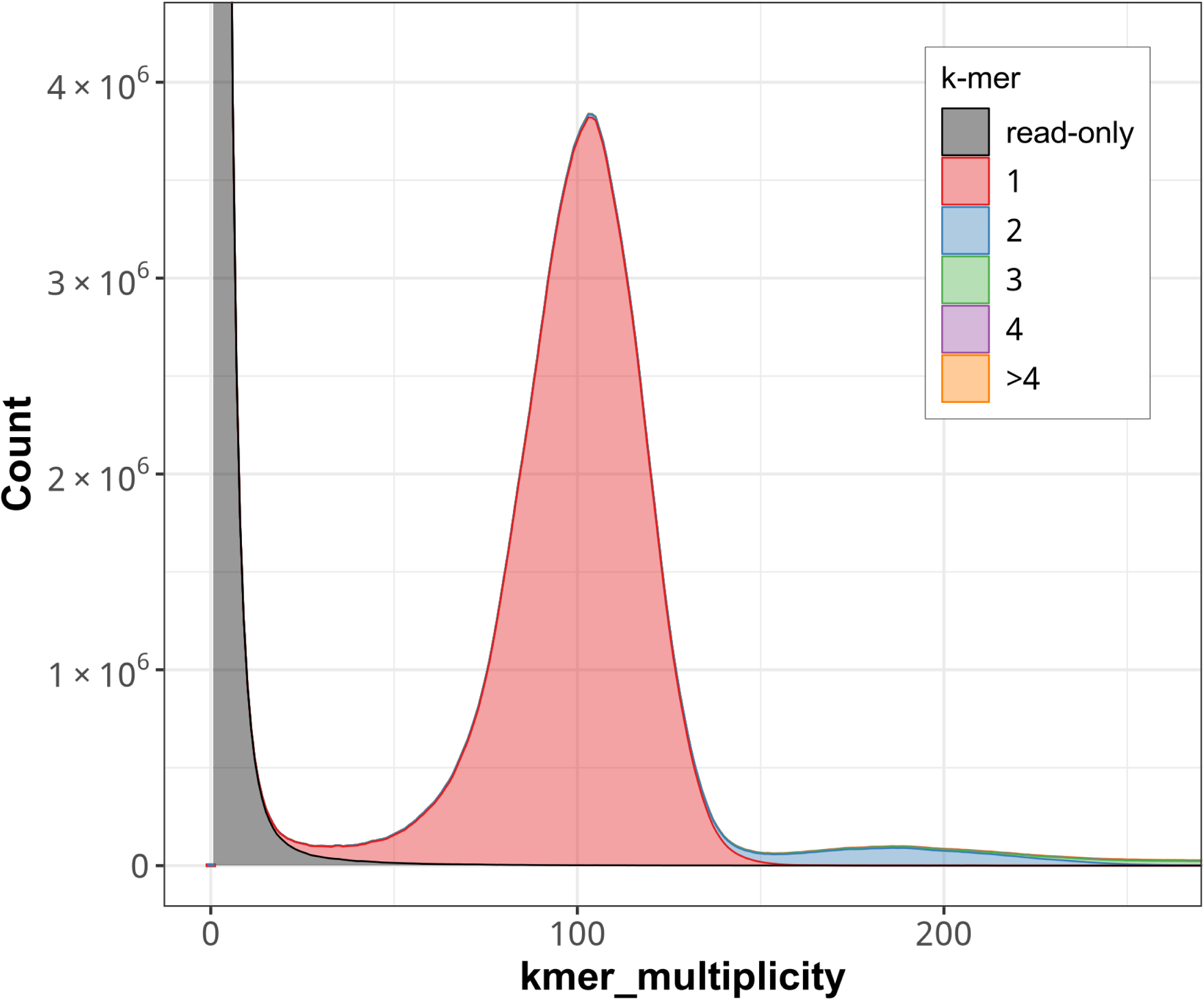
K-mer spectral analysis of the assembled genome data by Merqury. The graph was generated by Merqury software (version 1.3), with the default setting (k=21). The X axis shows read k-mer frequency; the Y axis shows number of k-mers.

**Table 3.** Results of BUSCO assessment

**Genome used as query**
| Category | Number of<br>identified BUSCOs | Percent of all BUSCO<br>database (insecta_odb10) |
| --- | --- | --- |
| Complete BUSCOs | 1353 | 99.0% |
| Complete and single-copy BUSCOs | 1349 | 98.7% |
| Complete and duplicated BUSCOs | 4 | 0.3% |
| Fragmented BUSCOs | 1 | 0.1% |
| Missing BUSCOs | 13 | 0.9% |
| Total | 1367 | 100% |

**Predicted proteina used as query**
| Category | Number of<br>identified BUSCOs | Percent of all BUSCO<br>database (insecta_odb10) |
| --- | --- | --- |
| Complete BUSCOs | 1339 | 98.0% |
| Complete and single-copy BUSCOs | 1230 | 90.0% |
| Complete and duplicated BUSCOs | 109 | 8.0% |
| Fragmented BUSCOs | 13 | 1.0% |
| Missing BUSCOs | 15 | 1.0% |
| Total | 1367 | 100% |

**Table 4.** Braconidae genome size available in NCBI Genome Data Viewer (Accessed in 2026/8/1)

| Species | Size | NCBI RefSeq or Genbank ID | Article DOI |
| --- | --- | --- | --- |
| Diachasmimorpha longicaudata | 194.5 Mb | GCF_034640455.1 |  |
| Diachasma alloeum | 384.4 Mb | GCF_001412515.2 | 10.1093/gbe/evz205 |
| Fopius arisanus | 153.6 Mb | GCF_000806365.1 | 10.1534/g3.117.040741 |
| Cotesia congregata | 199.2 Mb | GCA_905319865.3 |  |
| Cotesia typhae | 186.7 Mb | GCF_013202065.1 |  |
| Cotesia glomerata | 288.8 Mb | GCF_020080835.1 | 10.1093/jhered/esab032 |
| Microctonus hyperodae | 106.7 Mb | GCA_030347285.1 | 10.1186/s12864-023-09538-4 |
| Microctonus aethiopoides | 129.2 Mb | GCA_030347275.1 | 10.1186/s12864-023-09538-4 |
| Chelonus insularis | 135.7 Mb | GCF_013357705.1 | 10.1128/jvi.01573-21 |
| Microplitis deprimator | 233.2 Mb | GCA_964016945.1 | 10.12688/wellcomeopenres.23322.1 |
| Microplitis demolitor | 225.9 Mb | GCF_026212275.2 |  |
| Microplitis mediator | 269.3 Mb | GCF_029852145.1 |  |
| Aleiodes testaceus | 110.7 Mb | GCA_963565655.1 | 10.12688/wellcomeopenres.22738.1 |
| Aphidius gifuensis | 156.9 Mb | GCF_014905175.1 | 10.1111/1755-0998.13308 |
| *Ascogaster reticulata* (This study) | 280.5 Mb | - |  |

To assess the reliability of the predicted gene set, BUSCO analysis was performed using the predicted amino acid sequences as queries against the insecta_odb10 database. A total of 98% of BUSCOs were identified, of which 8% were classified as duplicated, whereas only 1% were classified as fragmented and 1% as missing (Table 3). Compared with the BUSCO assessment of the genome assembly, the predicted gene set contained a higher proportion of duplicated BUSCOs. This result is likely attributable to the gene-prediction process, which may have generated multiple gene models with highly similar sequences at the same or nearby genomic loci.

Despite the increased proportion of duplicated BUSCOs, the predicted gene set retained 98% of BUSCOs, indicating a high level of completeness. The number of predicted genes (27,261) was greater than those reported for other Braconidae species available in the NCBI Genome database (11,574 to 17,144; Table 5). This discrepancy is consistent with the elevated proportion of duplicated BUSCOs and suggests that some predicted genes may represent redundant gene models. Nevertheless, the high BUSCO completeness indicates that the gene set captures the vast majority of expected insect genes. Therefore, the predicted gene set should provide a comprehensive resource for gene-based analyses of *A. reticulata*.

**Table 5.** Numbers of genes in Braconidae species available in NCBI Genome Data Viewer (Accessed in 2026/8/1)

| Species | Number of Genes | NCBI RefSeq or Genbank ID | Article DOI |
| --- | --- | --- | --- |
| Diachasmimorpha longicaudata | 13,460 | GCF_034640455.1 |  |
| Diachasma alloeum | 13,480 | GCF_001412515.2 | 10.1093/gbe/evz205 |
| Fopius arisanus | 11,576 | GCF_000806365.1 | 10.1534/g3.117.040741 |
| Cotesia congregata | 14,119 | GCA_905319865.3 |  |
| Cotesia typhae | 14,743 | GCF_013202065.1 |  |
| Cotesia glomerata | 17,144 | GCF_020080835.1 | 10.1093/jhered/esab032 |
| Microctonus hyperodae | 11,688 | GCA_030347285.1 | 10.1186/s12864-023-09538-4 |
| Microctonus aethiopoides | 12,467 | GCA_030347275.1 | 10.1186/s12864-023-09538-4 |
| Chelonus insularis | 11,574 | GCF_013357705.1 | 10.1128/jvi.01573-21 |
| Microplitis deprimator | - | GCA_964016945.1 | 10.12688/wellcomeopenres.23322.1 |
| Microplitis demolitor | 13,141 | GCF_026212275.2 |  |
| Microplitis mediator | 15,924 | GCF_029852145.1 |  |
| Aleiodes testaceus | - | GCA_963565655.1 | 10.12688/wellcomeopenres.22738.1 |
| Aphidius gifuensis | 13,956 | GCF_014905175.1 | 10.1111/1755-0998.13308 |
| *Ascogaster reticulata* (This study) | 27,261 | - |  |

## Conclusion

The assembled genome and related data of *A. reticulata* reported in this study were found to be of sufficient quality as the reference genome data. Therefore, genome assembly, repeat annotation, predicted gene sets, and functional annotation datasets provide a genomic resource for comparative studies within Braconidae and for future investigations into the biology and evolution of Cheloninae parasitoid wasps.

## Acknowledgements

Parts of Figure 2 were drawn using illustrations from TogoTV (© 2016 DBCLS TogoTV, CC-BY-4.0 https://creativecommons.org/licenses/by/4.0/deed.ja ).

We would like to thank Editage (www.editage.jp) for English language editing.

## Author Contributions

K.Y., Y.K., K.K., and H.B. conceived the study. Y.K. collected samples. M.K. and S.K. prepared the genome DNA samples. K.Y. and H.B. performed the bioinformatic data analysis and data registration. K.Y., Y.K., S.K., K.K., and H.B. prepared original drafts of the manuscript. K.Y., Y.K., M.K., S.K., K.K., and H.B. reviewed and edited the manuscript. All the authors have read and agreed to the published version of the manuscript.

## Funding

This study was supported by the Center of Innovation for Bio-Digital Transformation (BioDX), an open innovation platform for industry-academia co-creation of the Japan Science and Technology Agency (JST) (COI-NEXT, grant number JPMJPF2010 to K.Y. and H.B.).

## Competing Interests

The authors declare that this study was conducted in the absence of competing interests.

## Notes

### Competing Interest Statement

The authors have declared no competing interest.

## References

1. Pennacchio, F. & Strand, M. R. Evolution of Developmental Strategies in Parasitic Hymenoptera. Annual Review of Entomology 51, 233–258 (2006).

2. Quicke, D. L. Parasitic Wasps. (Springer, London Weinheim, 1997).

3. Kawakami, T. Development of the Immature Stages of *Ascogaster reticulatus* WATANABE (Hymenoptera : Braconidae), an Egg-Larval : Parasitoid of the Smaller Tea Tortrix Moth, *Adoxophyes* sp.(Lepidoptera : Tortricidae). Applied Entomology and Zoology 20, 380–386 (1985).

4. Kainoh, Y. & Tamaki, Y. Searching Behavior and Oviposition of the Egg-Larval Parasitoid, *Ascogaster reticulatus* WATANABE (Hymenoptera : Braconidae). Applied Entomology and Zoology 17, 194–206 (1982).

5. Yoshiyama, M. et al. Possible Horizontal Transfer of a Transposable Element from Host to Parasitoid. Mol Biol Evol 18, 1952–1958 (2001).

6. Geib, S. M., Liang, G. H., Murphy, T. D. & Sim, S. B. Whole Genome Sequencing of the Braconid Parasitoid Wasp *Fopius arisanus*, an Important Biocontrol Agent of Pest Tepritid Fruit Flies. G3 Genes|Genomes|Genetics 7, 2407–2411 (2017).

7. Burke, G. R., Walden, K. K. O., Whitfield, J. B., Robertson, H. M. & Strand, M. R. Whole Genome Sequence of the Parasitoid Wasp *Microplitis demolitor* That Harbors an Endogenous Virus Mutualist. G3 Genes|Genomes|Genetics 8, 2875–2880 (2018).

8. Tvedte, E. S. et al. Genome of the Parasitoid Wasp *Diachasma alloeum*, an Emerging Model for Ecological Speciation and Transitions to Asexual Reproduction. Genome Biol Evol 11, 2767– 2773 (2019).

9. Beck, M. H., Zhang, S., Bitra, K., Burke, G. R. & Strand, M. R. The Encapsidated Genome of *Microplitis demolitor* Bracovirus Integrates into the Host Pseudoplusia includens. Journal of Virology 85, 11685–11696 (2011).

10. Drezen, J.-M., Chevignon, G., Louis, F. & Huguet, E. Origin and evolution of symbiotic viruses associated with parasitoid wasps. Current Opinion in Insect Science 6, 35–43 (2014).

11. Liu, J.-F., Zhao, H.-Y., Song, Y.-F., Yu, Y.-C. & Yang, M.-F. A Chromosome-Level Genome Assembly of the Parasitic Wasp *Chelonus formosanus* Sonan 1932 (Hymenoptera: Braconidae). Genome Biol Evol 14, evac006 (2022).

12. Kainoh, Y. Some Factors Influencing Sex Ratio in *Ascogaster reticulatus* WATANABE (Hymenoptera : Braconidae). Applied Entomology and Zoology 23, 35–40 (1988).

13. Tamaki, Y. Mass Rearing of the Smaller Tea Tortrix, *Adoxophyes orana* FISCHER VON ROSLERSTAMM, on a Simplified Artificial Diet for Successive Generations : (Lepidoptera : Tortricidae). Applied Entomology and Zoology 1, 120–124 (1966).

14. Cheng, H., Concepcion, G. T., Feng, X., Zhang, H. & Li, H. Haplotype-resolved de novo assembly using phased assembly graphs with hifiasm. Nat Methods 18, 170–175 (2021).

15. Cheng, H., Asri, M., Lucas, J., Koren, S. & Li, H. Scalable telomere-to-telomere assembly for diploid and polyploid genomes with double graph. Nat Methods 21, 967–970 (2024).

16. Cheng, H. et al. Haplotype-resolved assembly of diploid genomes without parental data. Nat Biotechnol 40, 1332–1335 (2022).

17. Assembly-stats. *Github* https://github.com/sanger-pathogens/assembly-stats.

18. Rhie, A., Walenz, B. P., Koren, S. & Phillippy, A. M. Merqury: reference-free quality, completeness, and phasing assessment for genome assemblies. Genome Biol 21, 245 (2020).

19. Manni, M., Berkeley, M. R., Seppey, M., Simão, F. A. & Zdobnov, E. M. BUSCO Update: Novel and Streamlined Workflows along with Broader and Deeper Phylogenetic Coverage for Scoring of Eukaryotic, Prokaryotic, and Viral Genomes. Molecular Biology and Evolution 38, 4647–4654 (2021).

20. Smit, A., Hubley, R. & Green, P. RepeatMasker Open-4.0.

21. Flynn, J. M. et al. RepeatModeler2 for automated genomic discovery of transposable element families. Proceedings of the National Academy of Sciences 117, 9451–9457 (2020).

22. Brůna, T., Hoff, K. J., Lomsadze, A., Stanke, M. & Borodovsky, M. BRAKER2: automatic eukaryotic genome annotation with GeneMark-EP+ and AUGUSTUS supported by a protein database. NAR Genomics and Bioinformatics 3, lqaa108 (2021).

23. Kriventseva, E. V. et al. OrthoDB v10: sampling the diversity of animal, plant, fungal, protist, bacterial and viral genomes for evolutionary and functional annotations of orthologs. Nucleic Acids Research 47, D807–D811 (2019).

24. NCBI genome Drosophila melanogaster https://www.ncbi.nlm.nih.gov/datasets/genome/GCF_000001215.4/. (2014).

25. NCBI genome Caenorhabditis elegans https://www.ncbi.nlm.nih.gov/datasets/genome/GCF_000002985.6/. (2013).

26. NCBI genome Homo sapiens https://www.ncbi.nlm.nih.gov/datasets/genome/GCF_000001405.40/. (2022).

27. NCBI genome Mus musculus https://www.ncbi.nlm.nih.gov/datasets/genome/GCF_000001635.27/. (2020).

28. The UniProt Consortium. UniProt: the Universal Protein Knowledgebase in 2025. Nucleic Acids Res 53, D609–D617 (2025).

29. Mistry, J. et al. Pfam: The protein families database in 2021. Nucleic Acids Research 49, D412–D419 (2021).

30. Bono, H., Sakamoto, T., Kasukawa, T. & Tabunoki, H. Systematic Functional Annotation Workflow for Insects. Insects 13, 586 (2022).

31. DNA Data Bank of Japan SRA https://ddbj.nig.ac.jp/search/entry/sra-submission/DRA029003 (2026).

32. NCBI Genbank https://www.ncbi.nlm.nih.gov/nuccore/BAAKUN01000000 (2026).

33. Yokoi, K., et al. List of contig IDs and Genbank IDs. Figshare 10.6084/m9.figshare.33095249 (2026).

34. Yokoi, K., et al. RepeatModeler and RepeatMasker output files. Figshare 10.6084/m9.figshare.33095321 (2026).

35. Yokoi, K., et al. Predicted gene set data. Figshare 10.6084/m9.figshare.33095726 (2026).

36. Yokoi, K., et al. Functional gene annotation data. Figshare 10.6084/m9.figshare.33095747 (2026).

37. Yokoi, K., et al. Supplemental data of ‘Genome sequence data of Ascogaster reticulata (Hymenoptera: Braconidae)’. 10.6084/m9.figshare.c.8618576 (2026).

38. Yokoi, K., et al. Script data. Figshare 10.6084/m9.figshare.33103445 (2026).

